# Airway Diameter-Matched Injury Improves Severity and Reproducibility of Experimental Rabbit Tracheal Stenosis

**DOI:** 10.64898/2026.07.12.738082

**Authors:** Benjamin M. Laitman, Claire Victoria Ong, Olivia Becker, Bryn Anderson, Grace W. Randall, David Gonzalez, Neha K. Reddy, Ya-Wen Chen

## Abstract

**Objective:** Reliable animal models of tracheal stenosis are necessary for the development and translational testing of anti-fibrotic and regenerative therapies, but existing rabbit models frequently demonstrate substantial variability in stenosis severity, which limits their translational utility. The objective of our study was to determine whether airway diameter-matched mechanical injury improves the severity and reproducibility of experimental tracheal stenosis in a rabbit model, and to evaluate whether rabbit body weight is a reliable surrogate for tracheal luminal diameter during model creation.

**Methods:** Fourteen male New Zealand White rabbits (weight range, 2.7–3.5 kg) underwent tracheal injury using steel-bristle brushes introduced through a tracheotomy. Animals were assigned to receive either airway diameter-matched injury, in which brush size was selected to closely approximate the directly measured tracheal lumen diameter, or non-matched injury, in which brush size was selected without regard to measured lumen diameter. At postoperative day 21 (POD21), the injured tracheal segment and a native uninjured segment from the same animal were harvested and compared. Stenosis degree was quantified grossly, and lamina propria-to-cartilage (LP:C) ratio was quantified histologically by three blinded reviewers. The relationship between rabbit weight and airway diameter was assessed, and inter-rater reliability was calculated using the intraclass correlation coefficient (ICC).

**Results:** Twelve of fourteen rabbits reached the POD21 endpoint; two were euthanized early for severe airway compromise meeting humane endpoint criteria, both with approximately 80% stenosis. Injured tracheas demonstrated significantly greater stenosis than native controls (66.0 ± 13.0% vs 16.0 ± 2.7%; p = 0.00012), with a corresponding increase in LP:C ratio (p = 0.031). Airway diameter-matched injury produced significantly greater stenosis than non-matched injury (74.6 ± 6.1% vs 50.6 ± 4.0%; p = 0.001), while LP:C ratio did not differ between injury techniques (p = 1.0). Rabbit weight did not correlate with airway diameter (r = 0.176, p = 0.515; R^2^ = 0.031). Inter-rater reliability was excellent for both stenosis degree (ICC = 0.989) and LP:C ratio (ICC = 0.992).

**Conclusions:** Direct measurement and matching of injury instrument diameter to native airway diameter substantially improves both the severity and the reproducibility of stenosis in a rabbit tracheal injury model, whereas body weight is an unreliable surrogate for airway size. This optimized, standardized protocol offers a reproducible platform for future translational studies of airway fibrosis and anti-fibrotic or regenerative therapies.

## INTRODUCTION

Laryngotracheal stenosis is a debilitating condition that arises from a host of etiologies including prolonged endotracheal intubation, tracheostomy, external airway trauma, autoimmune disease, or infection.^1,2^ Affected patients commonly experience dyspnea, reduced exercise tolerance, and the need for recurrent endoscopic or open procedures, all of which can meaningfully impair quality of life.^3^ Long-segment stenosis is particularly difficult to manage definitively, and at present no therapy reliably prevents the fibrotic scarring that follows airway injury.^4,5^

Development of anti-fibrotic and regenerative strategies for airway stenosis depends on reproducible preclinical models in which injury severity can be controlled and outcomes reliably measured. The rabbit has long served as a workhorse species for this purpose: its airway is surgically accessible, its dimensions are appropriate for instrumented injury and endoscopic follow-up, its tissue is amenable to detailed histologic evaluation, and the model is comparatively cost-effective relative to larger animal systems.^6, 7^ Despite these advantages, numerous rabbit tracheal stenosis models described in the literature report substantial variability in the degree of stenosis achieved, both within and across cohorts, and reproducibility between investigators and institutions has remained a persistent limitation.^8, 9, 10^ Most published protocols apply a single, fixed-diameter injury instrument to all animals within a broad weight range (commonly 2.5–3.5 kg), implicitly assuming that body weight is a reasonable proxy for tracheal luminal diameter.^11,12^ Whether this assumption is valid has not, to our knowledge, been systematically tested.

We hypothesized that fixed-instrument injury protocols produce variable stenosis because airway diameter varies independently of body weight, and that matching the injury instrument to the directly measured native airway diameter of each animal would produce more severe and more reproducible stenosis. The objective of this study was therefore to determine whether airway diameter-matched mechanical injury improves the severity and reproducibility of experimental rabbit tracheal stenosis relative to non-matched injury, and to establish a standardized, generalizable protocol for future translational investigation.

## MATERIALS AND METHODS

### Animal Subjects

This prospective, controlled animal study was approved by our Institutional Animal Care and Use Committee (IACUC) prior to initiation. Fourteen male New Zealand White rabbits weighing 2.7– 3.5 kg were enrolled and housed under daily veterinary observation throughout the study period. Animal subjects were divided into 2 experimental groups: 8 rabbits in a non-size matched group and 6 rabbits in a size-matched experimental group. Characteristics of each rabbit in both groups is summarized in **Table 1**.

**Table 1.** Summary of rabbit label, experimental group, weight, trachea lumen diameter and diameter of brush used during surgery.

| Rabbit ID | Experimental Group | Weight (kg) | Trachea Lumen Diameter (mm) | Brush Diameter (mm) |
| --- | --- | --- | --- | --- |
| R1 | Non-matched injury | 3.1 | 6.6 | 4.5 |
| R2 | Non-matched injury | 3.26 | 5.9 | 5.5 |
| R3 | Non-matched injury | 3.4 | 6.3 | 4.5 |
| R4 | Non-matched injury | 3.15 | 6.2 | 5.5 |
| R5 | Non-matched injury | 3 | 6.3 | 5.5 |
| R6 | Non-matched injury | 3.9 | 5.4 | 4.5 |
| R7 | Non-matched injury | 3.3 | 6.6 | 3.5 |
| R8 | Non-matched injury | 3.4 | 6.5 | 4.5 |
| R9 | Matched injury | 2.7 | 5 | 4.5 |
| R10 | Matched injury | 3.26 | 6 | 5 |
| R11 | Matched injury | 3 | 6 | 5.5 |
| R12 | Matched injury | 2.9 | 6 | 5 |
| R13 | Matched injury | 3 | 6 | 5.5 |
| R14 | Matched injury | 3.3 | 6 | 5.5 |

### Surgical Procedure

All procedures were performed by authors CVO, DG, or BML. General anesthesia was induced with isoflurane, and animals underwent endotracheal intubation. The planned surgical field was locally infiltrated with 2% lidocaine hydrochloride, and a midline cervical incision was made. After division of the platysma and blunt separation of the sternohyoid and sternothyroid muscles, the trachea was exposed approximately 3.5 cm inferior to the cricothyroid ligament. The endotracheal tube was retracted. A tracheal incision encompassing approximately two-thirds of the tracheal circumference was then created, and the trachea was gently elevated until the lumen assumed a circular configuration, at which point the airway diameter was measured directly.

Animals were assigned to receive either airway diameter-matched or non-matched mechanical injury. In the diameter-matched group, the injury brush was selected to closely approximate the directly measured tracheal lumen diameter within ∼0.5 mm (for example, a 6.0 mm airway received a 5.5 mm brush), whereas in the non-matched group, brush size was selected without regard to the measured lumen diameter. Two of the rabbits in the circumference matched group had brushes 1mm below the tracheal lumen diameter as 0.25 mm brush adjustments were not feasible. Circumferential mucosal injury was then performed using steel-bristle brushes (MSC Industrial Supply Co., Melville, NY) available in diameters of 3.5, 4.0, 4.5, 5.0, 5.5, 6.0, 6.5, and 7.0 mm. Brushes were inserted 1 cm into the lumen and ten rotational passes applied to the tracheal mucosa in each animal. A pediatric suction catheter was passed proximally and distally to clear blood and debris from the airway, hemostasis was achieved with epinephrine-soaked cotton, and the trachea was closed with 6-0 polypropylene suture. The muscle and subcutaneous tissue were closed with 3-0 Vicryl suture, and the skin was closed with 3-0 nylon suture.

Animals were observed daily by veterinary staff, with behavioral assessments recorded and body weight measured on the day of surgery (POD0) and on postoperative days 2, 4, 7, 14, and 21. Predefined humane endpoint criteria included respiratory distress, hypoxemia, reduced activity, and failure to thrive. Steps of surgical procedure outlined in **Figure 1**.

**Figure 1.**
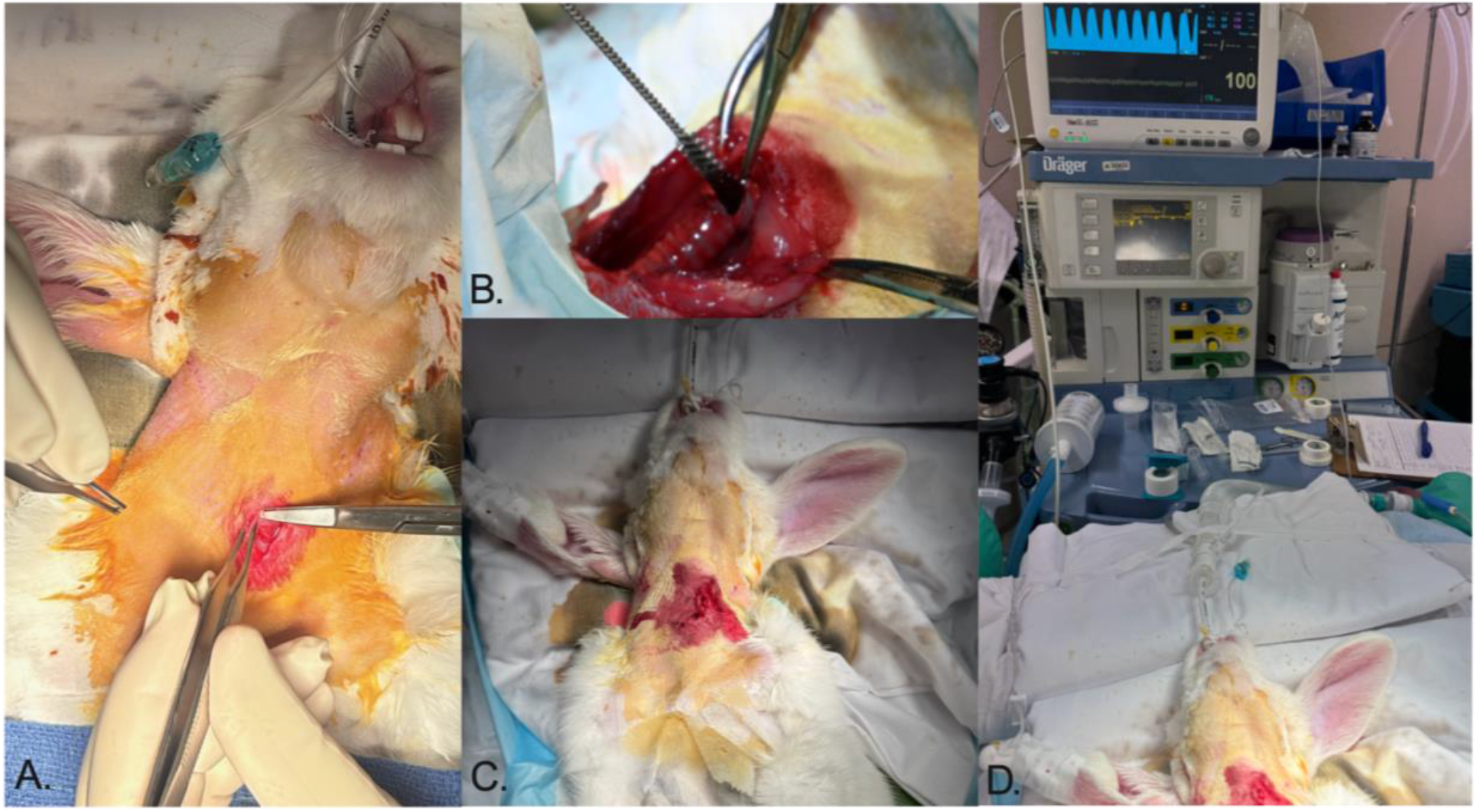
Surgical procedure. **(A)** Following endotracheal intubation and induction of anesthesia, a midline cervical incision was performed and platysma divided to expose the underlying neck strap muscles and trachea. **(B)** A steel-bristle brush was inserted into the tracheal lumen and rotated circumferentially 10 times to induce mucosal injury. **(C)** Following injury the trachea was closed, and the neck strap muscles, platysma, and skin were reapproximated in layers. **(D)** Animals were continuously monitored during surgery and recovery using electrocardiography, pulse oximetry, and respiratory waveform monitoring to assess cardiac and respiratory status.

### Specimen Collection and Preparation

At POD21, the injured tracheal segment and a native, uninjured tracheal segment were harvested from each animal; using each rabbit as its own internal control minimized confounding related to inter-animal variability in baseline airway size. POD21 was chosen as the primary endpoint as this has been shown to be a time of maximal airway stenosis in prior literature (REFS). Percent stenosis was calculated for each segment according to the method of Chen et al. (2024) ^12^, using the longest and shortest luminal diameters (*rl* and *rs*, respectively) and the longest and shortest tracheal diameters (*RL* and *RS*, respectively).

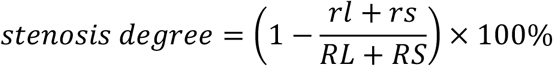

Tracheal segments were subsequently fixed in formalin, paraffin-embedded, sectioned at 5 µm, and stained with hematoxylin and eosin according to the protocol provided in the supplementary methods. Sections were evaluated for epithelial morphology, lamina propria expansion, subepithelial fibrosis, and cartilage architecture. At three locations around the tracheal circumference, lamina propria thickness (from the basement membrane to the inner cartilage border) and cartilage thickness (from the inner to the outer cartilage border) were measured, and the average measurements at each location were used to calculate the lamina propria-to-cartilage (LP:C) ratio. All gross and histologic measurements were performed independently by three blinded reviewers, and reproducibility across reviewers was assessed using the intraclass correlation coefficient (ICC). Gross anatomy and histologic samples of native and stenosed trachea along with example of stenosis measurements depicted in **Figure 2**.

**Figure 2.**
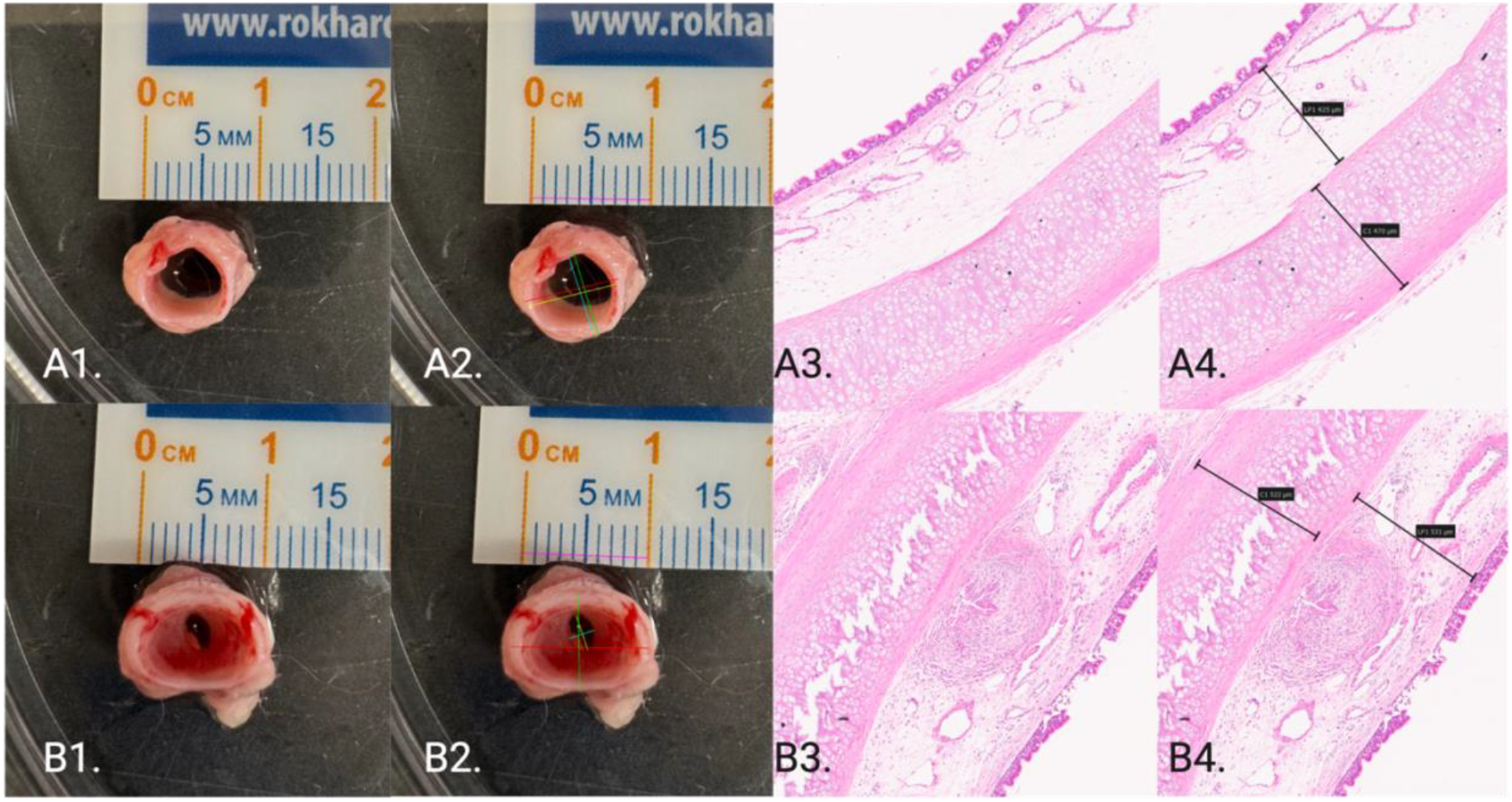
Gross and histologic quantification of experimental tracheal stenosis. Representative native trachea (**A**) and stenosed trachea (**B**) harvested at postoperative day 21. (**A1, B1**) Gross tracheal specimens following harvest. (**A2, B2**) Gross stenosis measurements performed using Fiji/ImageJ. The magenta line represents the scale reference, yellow denotes the longest luminal diameter (*rl*), cyan denotes the shortest luminal diameter (*rs*), red denotes the longest tracheal diameter (*RL*), and green denotes the shortest tracheal diameter (*RS*). Stenosis degree was calculated according to the method described by Chen et al.^12^ (**A3, B3**) Representative hematoxylin and eosin (H&E)-stained sections of native and stenosed trachea. (**A4, B4**) Histologic measurements used to calculate the lamina propria-to-cartilage (LP:C) ratio. Lamina propria thickness (LP) was measured from the epithelial basement membrane to the inner cartilage border, while cartilage thickness (C) was measured from the inner to outer cartilage border. Measurements were obtained at three locations distributed around the tracheal circumference and averaged for analysis.

### Statistical Analysis

Given the modest sample size and non-normal data distribution, nonparametric statistical methods were used throughout. Stenosis degree and LP:C ratio were compared between native and stenosed segments using the Wilcoxon signed-rank test, and between airway-matched and non-matched injury groups using the Wilcoxon rank-sum test. The relationship between rabbit weight and airway diameter was assessed using Pearson correlation and linear regression (R^2^), and inter-rater reliability was assessed using a two-way, average-measures intraclass correlation coefficient. All statistical analyses were performed in RStudio (RStudio Team, Boston, MA), and a p-value less than 0.05 was considered statistically significant. With sample sizes of 8 (non-matched) and 6 (matched injury), this study had 80% power (α = 0.05) to detect a minimum absolute difference of approximately 17 percentage points in gross stenosis between groups.

## RESULTS

Summary of stenosis analysis results per group is summarized in **Table 2**.

**Table 2.**
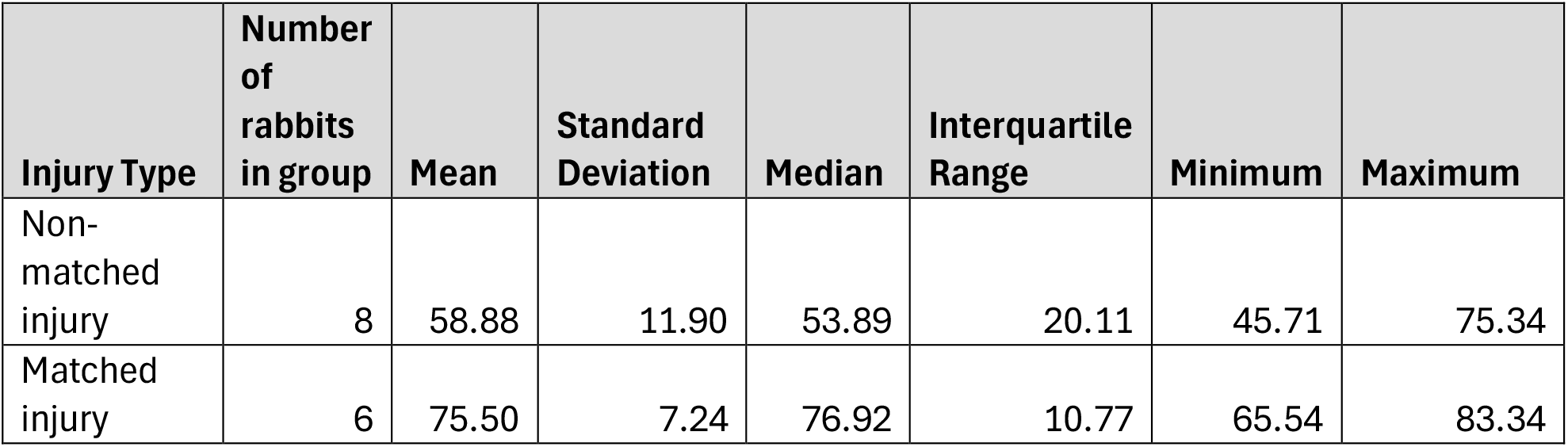
Summary of descriptive statistics for non-matched vs matched injury experimental groups.

### Animal Outcomes

Fourteen rabbits were enrolled, and twelve reached the POD21 study endpoint. Two rabbits were euthanized prior to the planned endpoint (one at POD19 and the other POD20) because of severe airway compromise meeting predefined humane endpoint criteria; both animals demonstrated approximately 80% stenosis at the time of euthanasia. No animal developed postoperative infection, bleeding, or wound complications. Mild weight loss and transient reductions in activity were observed during POD2–4, with recovery of both weight and activity by approximately POD7. (**Supplementary Figure A**).

### Mechanical Injury Produces Significant Tracheal Stenosis

Inter-rater reliability was excellent for both gross and histologic outcome measures. For stenosis degree, ICC was 0.989 (95% CI, 0.954–0.996). For LP:C ratio, ICC was 0.992 (95% CI, 0.977– 0.998).

Native tracheal segments demonstrated a mean luminal reduction of 16.0 ± 2.7%, compared with ± 13.0% in stenosed segments from the same animals (Wilcoxon signed-rank, p = 0.00012), regardless of brush matching **(Figure 3)**. The LP:C ratio was significantly increased in stenosed segments compared with native controls (Wilcoxon signed-rank, p = 0.031) (**Figure 4**).

**Figure 3.**
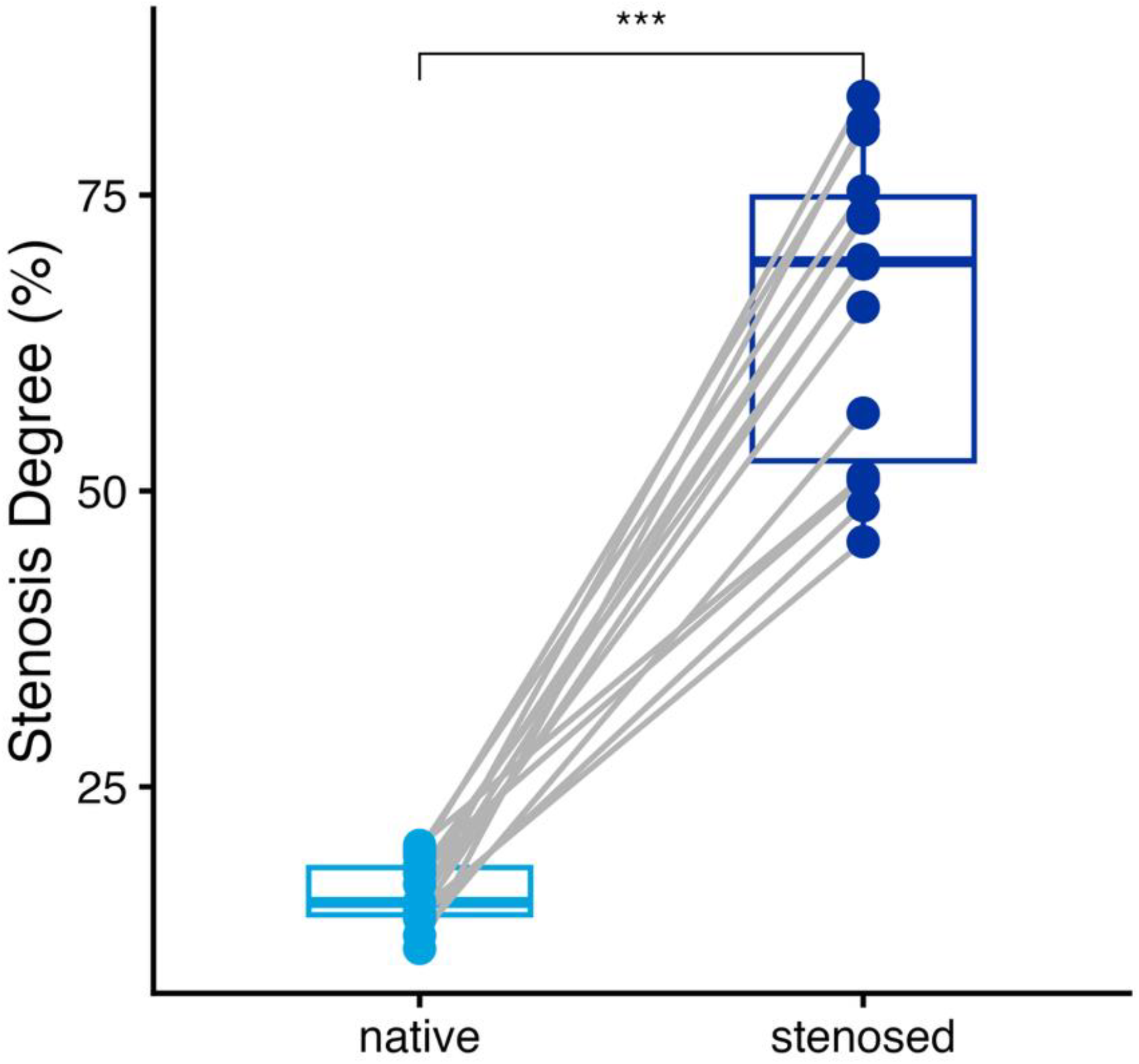
Percent stenosis in native versus mechanically injured tracheal segments at POD21 (66.0 ± 13.0% vs 16.0 ± 2.7%; p = 0.00012).

**Figure 4.**
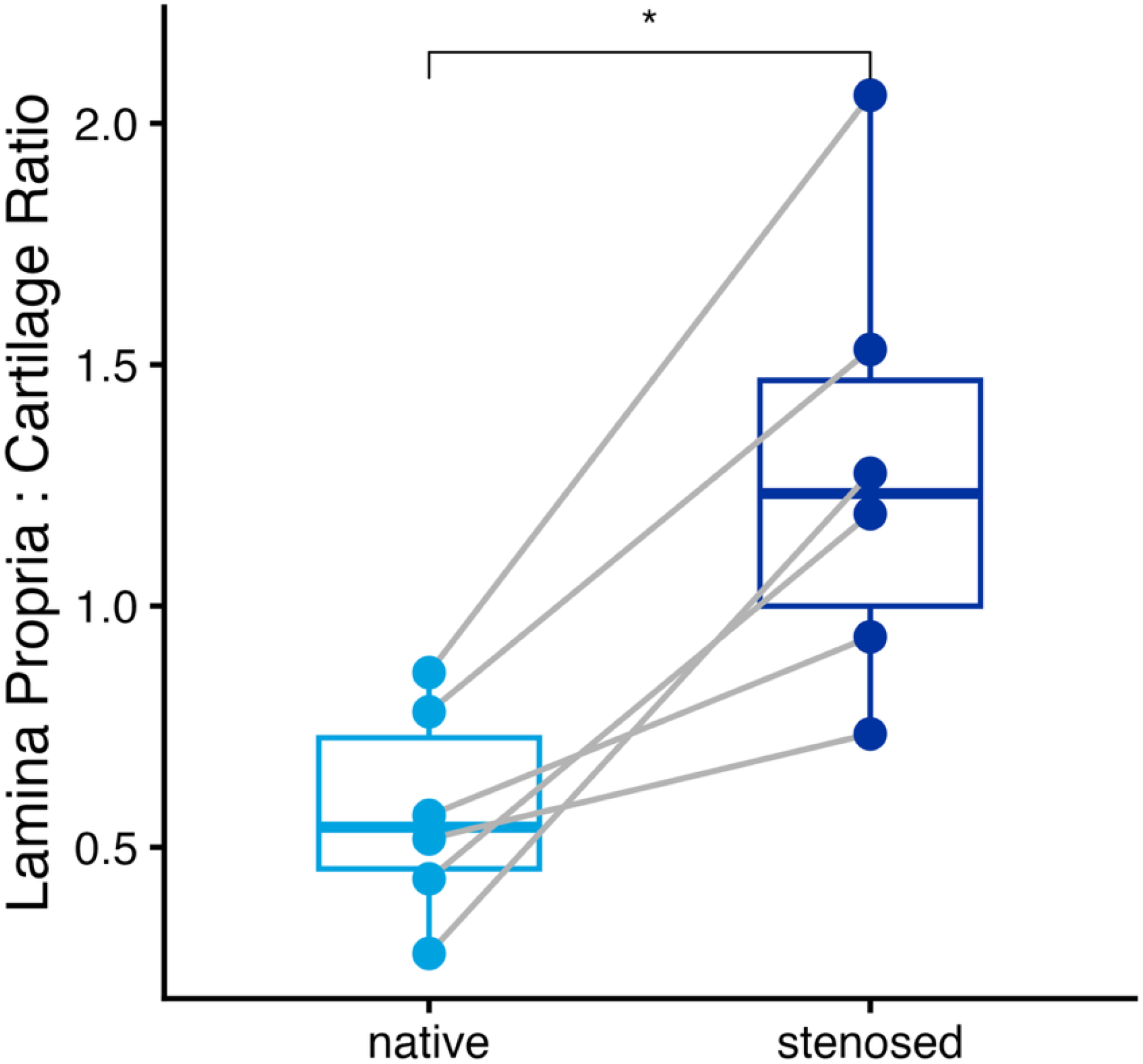
Lamina propria-to-cartilage (LP:C) ratio in native versus stenosed tracheal segments (p = 0.031).

### Airway Diameter-Matched Injury Produces Greater Stenosis and Less Variability Between Injury Attempt

Non-matched injury produced a mean stenosis of 58.9 ± 11.9% (median 53.9%, IQR 20.1%), compared with 75.5 ± 7.2% (median 76.9%, IQR 10.8%) with airway diameter-matched injury (Wilcoxon rank-sum, p = 0.001) (**Figure 5**).

**Figure 5.**
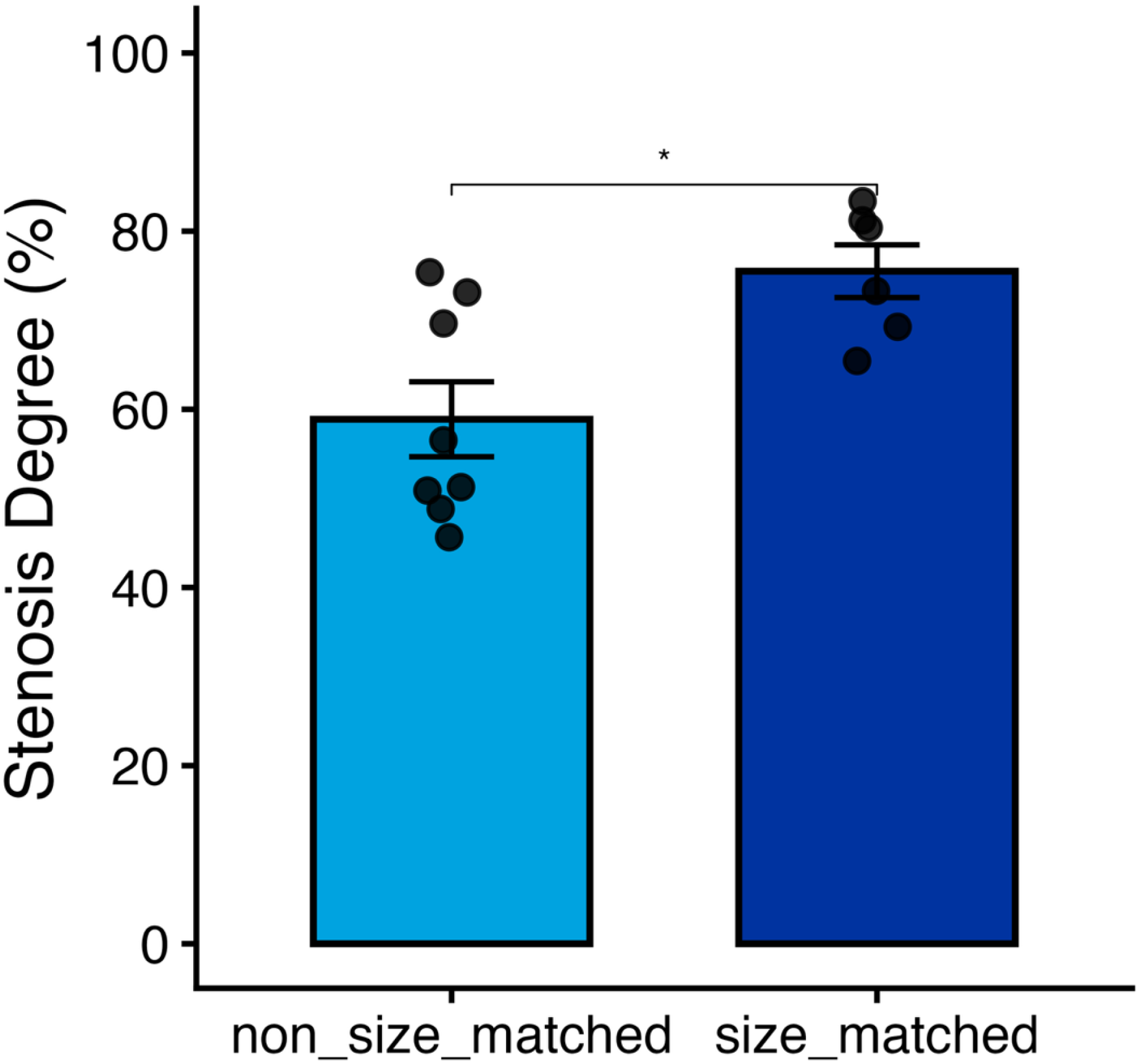
Percent stenosis in airway diameter-matched versus non-matched injury groups (74.6 ± 6.1% vs 50.6 ± 4.0%; p = 0.001).

### Rabbit Weight Does Not Predict Airway Diameter

Rabbit weight did not correlate with tracheal airway diameter (Pearson r = 0.176, p = 0.515). Linear regression demonstrated an R^2^ of 0.031, indicating that body weight explained only 3.1% of the variability in airway diameter (**Figure 6**).

**Figure 6.**
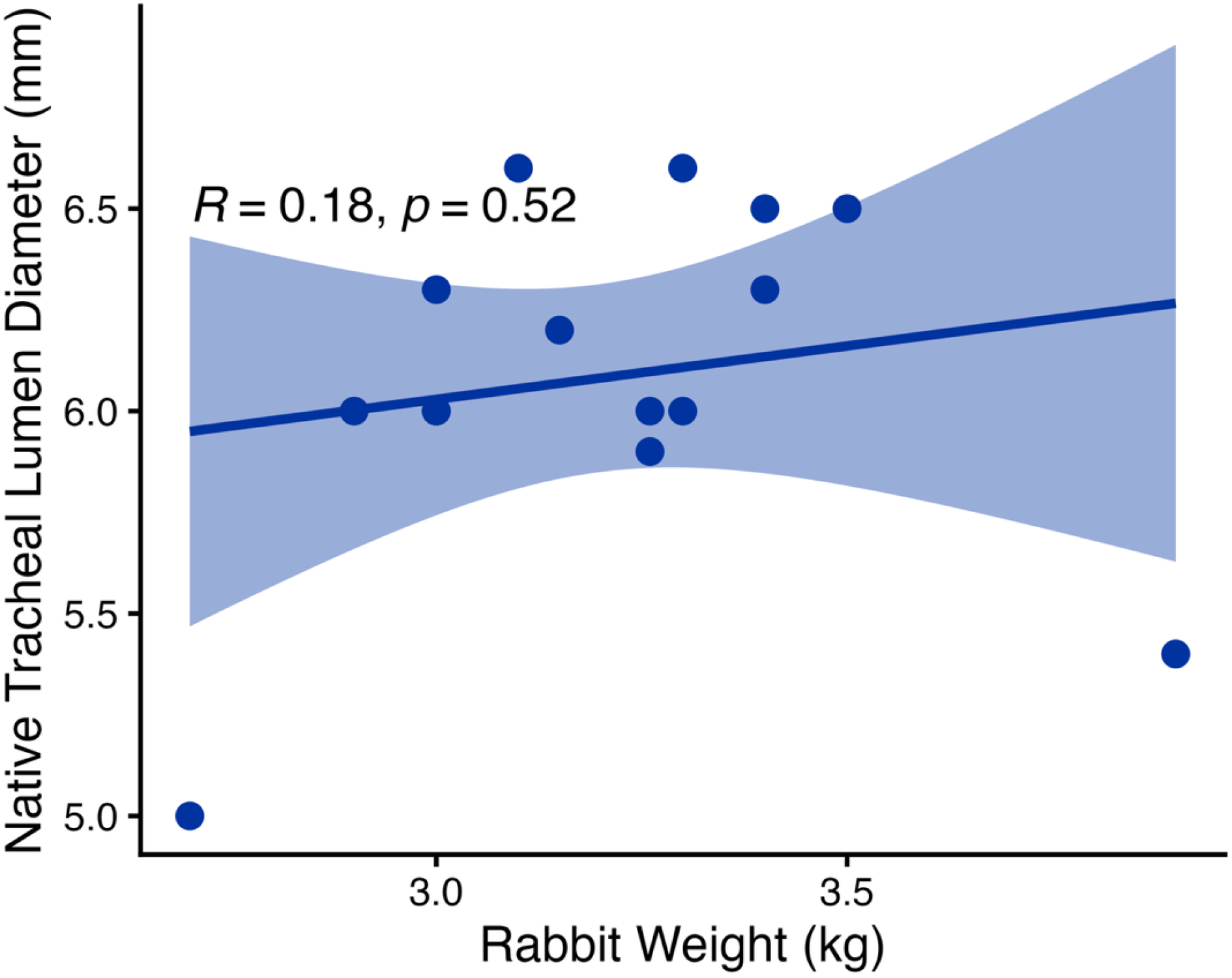
Correlation between rabbit weight and tracheal airway diameter (r = 0.176, R^2^ = 0.031).

## DISCUSSION

Circumferential mechanical injury with a steel brush produced significant gross stenosis and histologic remodeling in every animal, with consistent stenosis achieved across the cohort, confirming what has been previously reported in the literature: that brush-based mucosal injury is an effective means of generating a fibrotic stenosis phenotype in the rabbit trachea.^7,9,12^ In the non-matched injury group, the mean stenosis was 58.9%, which agrees with prior nylon-brush models, which have reported highly variable results, ranging from 10% to 80%,^8,10,11^ In contrast, steel brush injury has been shown to induce more reliable stenosis than nylon brush techniques,^13^ and present findings are consistent with this observation.

Building on this baseline finding, matching the injury instrument to each animal’s directly measured airway diameter increased stenosis severity by approximately 24 percentage points relative to non-matched injury (mean stenosis of 75.5% in airway-matched injury group). This finding indicates that individualized injury selection, rather than reliance on a fixed instrument size, is necessary to achieve maximal and consistent luminal narrowing, and suggests that matching injury to native airway anatomy is a simple technical modification that substantially improves the performance of the model. Reproducibility remains one of the principal challenges in experimental tracheal stenosis research. By standardizing injury magnitude relative to each animal’s native airway rather than to a fixed absolute instrument size, diameter-matched injury produced more predictable and tightly distributed stenosis formation, as reflected in the narrower variance observed in the matched group. This standardized approach should improve reproducibility not only within a single laboratory but also between investigators and institutions.

These findings also help explain why prior rabbit tracheal stenosis models have demonstrated such variable results. Most published protocols apply a single injury instrument size across all animals within a broad weight range, implicitly assuming that body weight is a reasonable proxy for airway size. However, the present study demonstrates that rabbit weight does not reliably predict airway diameter, such that two rabbits of similar weight may have substantially different tracheal luminal dimensions. Consequently, an identical injury instrument applied across a weight-defined cohort may produce markedly different injury severities from animal to animal, as a varying degree of lumen may be contacted by the brush diameter. This underscores the importance of direct airway measurement during model creation. Further, our use of each rabbit as its own control, by harvesting both injured and native uninjured segment of trachea from the same animal, allows for a more precise measurement of stenosis using a more accurate control measurement of luminal area.

Taken together, this work establishes a standardized, reproducible rabbit model of tracheal stenosis that is well suited to the evaluation of anti-fibrotic therapies, local drug delivery systems, biomaterials, tissue engineering constructs, and other regenerative medicine strategies aimed at preventing or reversing airway fibrosis. Several limitations should nonetheless be considered when interpreting these findings. The sample size was modest, and outcomes were assessed at a single postoperative timepoint (POD21) without serial bronchoscopic or radiographic imaging or intermediate histologic evaluation. As an acute mechanical injury model, this protocol does not fully recapitulate the pathophysiology of stenosis following prolonged endotracheal intubation or autoimmune conditions. Finally, the study was limited to a single species and a single injury modality, and further work is needed to determine whether these findings generalize across other injury mechanisms and animal models.

## CONCLUSION

Airway diameter-matched mechanical injury produces significantly greater and more reproducible tracheal stenosis than non-matched injury in a rabbit model. Rabbit body weight does not predict airway diameter, supporting the use of direct airway measurement, rather than weight-based instrument selection, during model creation. This optimized and standardized protocol provides a reproducible platform for future translational investigations of airway fibrosis, anti-fibrotic therapies, and airway regeneration.

## ACKNOWLEDGMENTS

We would like to thank the Icahn School of Medicine Center for Comparative Medicine and Surgery for their help with caring for our rabbits.

---

**Supplementary Figure A.**
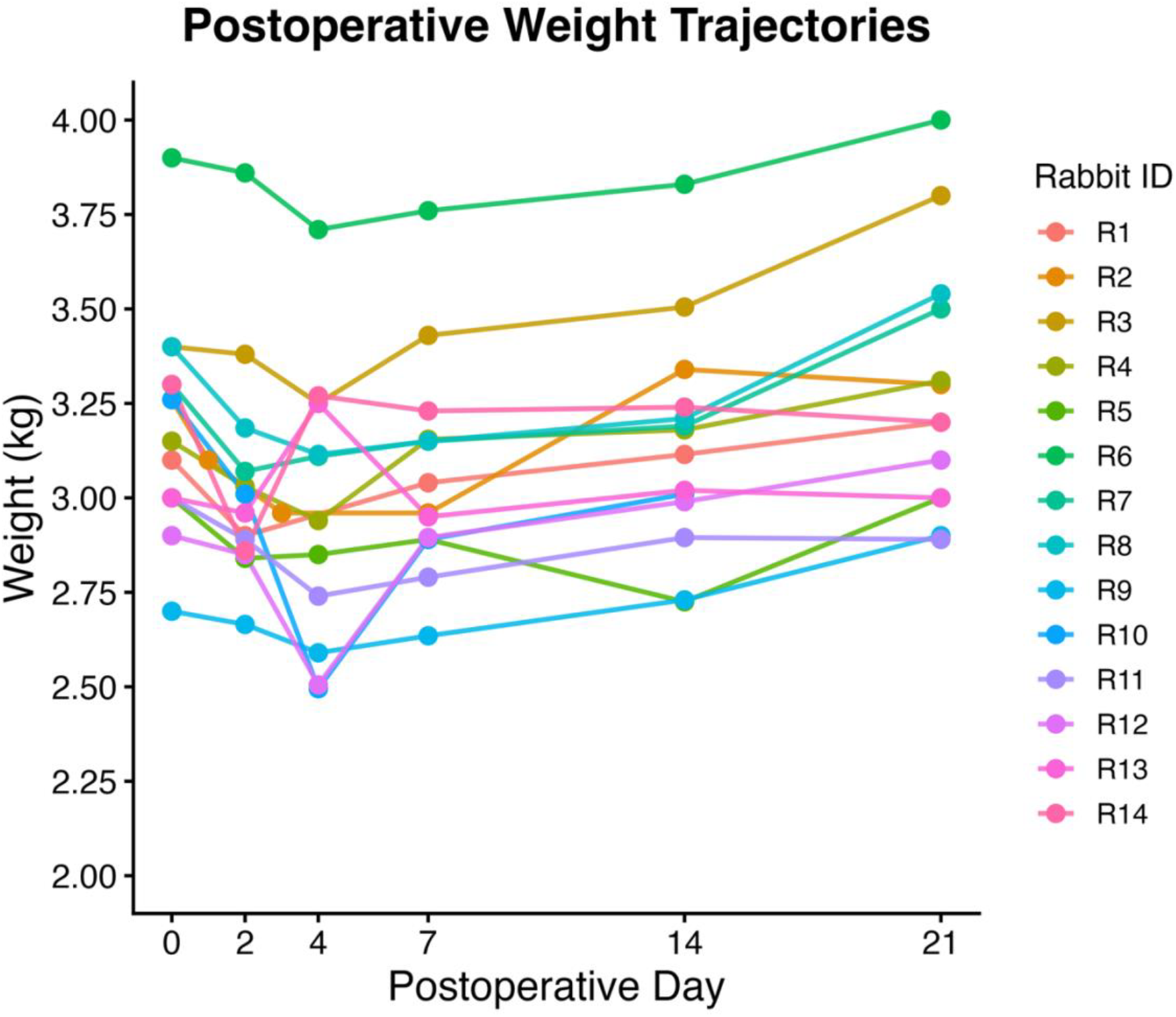
Postoperative weight trajectory (POD0–POD21).

## REFERENCES

1. Carpenter, D. J., Hamdi, O. A., Finberg, A. M., & Daniero, J. J. (2022). Laryngotracheal stenosis: Mechanistic review. Head & neck, 44(8), 1948–1960. 10.1002/hed.27079.

2. Curry, S. D., & Rowan, P. J. (2020). Laryngotracheal Stenosis in Early vs Late Tracheostomy: A Systematic Review. Otolaryngology--head and neck surgery : official journal of American Academy of Otolaryngology-Head and Neck Surgery, 162(2), 160–167. 10.1177/0194599819889690.

3. Smith, M. M., & Cotton, R. T. (2018). Diagnosis and management of laryngotracheal stenosis. Expert review of respiratory medicine, 12(8), 709–717. 10.1080/17476348.2018.1495564.

4. Perryman, M. C., Kraft, S. M., & Kavookjian, H. L. (2023). Laryngotracheal Reconstruction for Subglottic and Tracheal Stenosis. Otolaryngologic clinics of North America, 56(4), 769–778. 10.1016/j.otc.2023.04.018.

5. Wen, W., Du, X., Zhu, L., Wang, S., Xu, Z., & Lu, Z. (2022). Surgical management of long-segment congenital tracheal stenosis with tracheobronchial malacia. European journal of cardio-thoracic surgery : official journal of the European Association for Cardio-thoracic Surgery, 61(5), 1001–1010. 10.1093/ejcts/ezab551

6. Lee, H. S., Kim, S. W., Oak, C., Ahn, Y. C., Kang, H. W., Chun, B. K., & Lee, K. D. (2015). Rabbit model of tracheal stenosis induced by prolonged endotracheal intubation using a segmented tube. International journal of pediatric otorhinolaryngology, 79(12), 2384–2388. 10.1016/j.ijporl.2015.10.049.

7. Nakagishi, Y., Morimoto, Y., Fujita, M., Ozeki, Y., Maehara, T. and Kikuchi, M. (2005), Rabbit Model of Airway Stenosis Induced by Scraping of the Tracheal Mucosa. The Laryngoscope, 115: 1087–1092. 10.1097/01.MLG.0000163105.86513.6D.

8. Steehler, M.K., Hesham, H.N., Wycherly, B.J., Burke, K.M. and Malekzadeh, S. (2011), Induction of tracheal stenosis in a rabbit model—endoscopic versus open technique. The Laryngoscope, 121: 509–514. 10.1002/lary.21407.

9. McIlwain, W. R., Wistermayer, P. R., Swiss, T. P., Marko, S. T., Ieronimakis, N. M., & Rogers, D. J. (2017). Reproducing severe acute subglottic stenosis in a rabbit model. International journal of pediatric otorhinolaryngology, 103, 142–146. 10.1016/j.ijporl.2017.10.011.

10. Schweiger, C., Hart, C. K., Tabangin, M. E., Cohen, A. P., Roetting, N. J., DeMarcantonio, M., Becker, E., Ward, J. A., & de Alarcón, A. (2019). Development of a survival animal model for subglottic stenosis. The Laryngoscope, 129(4), 989–994. 10.1002/lary.27441.

11. Zhang, J., Liu, Y. H., Yang, Z. Y., Liu, Z. Y., Wang, C. G., Zeng, D. X., & Jiang, J. H. (2023). The role of tracheal wall injury in the development of benign airway stenosis in rabbits. Scientific reports, 13(1), 3144. 10.1038/s41598-023-29483-2

12. Chen, W., Wang, Q., Xu, H., Xie, Y., Zhang, L., Li, Y., Yan, G., Ding, Y., Lu, S., Xie, Z., Chen, J., Xu, M., Liang, X., Chen, J., Fu, P., Li, X., & Peng, L. (2024). Establishment of a survival rabbit model for laryngotracheal stenosis: A prospective randomized study. Laryngoscope investigative otolaryngology, 9(6), e70047. 10.1002/lio2.70047.

13. Lin, H., Ainiwaer, M., Jiang, Z., Wang, Z., Liu, J., & Chen, F. (2024). Comparative evaluation of mechanical injury methods for establishing stable tracheal stenosis animal models. Scientific reports, 14(1), 2383. 10.1038/s41598-024-52230-0.

